# LM-GVP: A Generalizable Deep Learning Framework for Protein Property Prediction from Sequence and Structure

**DOI:** 10.1101/2021.09.21.460852

**Authors:** Zichen Wang, Steven A. Combs, Ryan Brand, Miguel Romero Calvo, Panpan Xu, George Price, Nataliya Golovach, Emmanuel O. Salawu, Colby J. Wise, Sri Priya Ponnapalli, Peter M. Clark

## Abstract

Proteins perform many essential functions in biological systems and can be successfully developed as bio-therapeutics. It is invaluable to be able to predict their properties based on a proposed sequence and structure. In this study, we developed a novel generalizable deep learning framework, LM-GVP, composed of a protein Language Model (LM) and Graph Neural Network (GNN) to leverage information from both 1D amino acid sequences and 3D structures of proteins. Our approach outperformed the state-of-the-art protein LMs on a variety of property prediction tasks including fluorescence, protease stability, and protein functions from Gene Ontology (GO). We also illustrated insights into how a GNN prediction head can guide the protein LM to better leverage structural information. We envision that our deep learning framework will be generalizable to many protein property prediction problems to greatly accelerate protein engineering and drug development.

## Introduction

Proteins are the major macromolecules carrying out the essential functions in biology. Composed of a sequence of amino acids (AAs) connected by peptide bonds, a natural protein folds into its tertiary (3D) structure during biosynthesis on the ribosome^1^ to carry out its function. Artificially designed proteins or polypeptide chains can also be synthesized to exhibit desired biological functions for research and therapeutic applications.

An important problem researchers have been studying intensively for over five decades is the complete determination of 3D protein structures, which ultimately enhances our understanding of many of their other properties such as biological function, druggability, and stability against physical or enzymatic stress. Protein structures can be determined experimentally using methods such as nuclear magnetic resonance (NMR), X-ray crystallography, and cryogenic electron microscopy. Computational methods for predicting unknown structures have also been developed via software like Rosetta. Recent breakthroughs in deep learning approaches including AlphaFold^2,3^, trRosetta^4^ and a three-track deep neural net approach by Baek et al.^5^, have exceeded performance based on traditional approaches for modeling the folding processes.

However, accurately predicting the 3D structures is merely the first step in protein modeling. The ultimate goal is to predict proteins’ properties. The AlphaFold2^3^ study has demonstrated that sequence alone can predict 3D structures with outstanding performance, thus is it still necessary to explicitly incorporate known structural information into protein models? Just as phenotypes are not fully determined by genotype, so too are protein sequences not always folded into the same 3D structures when subjected to different physiological conditions. An extreme example is the mis-folded forms of Amyloid beta, which are implicated in Alzheimer’s disease^6^. The conformation of proteins’ 3D structures, such as G-protein-coupled receptors (GPCRs)^7^ and hemoglobin, also change with the presence or absence of allosteric modulators, which directly affects protein functions such as ligand binding and oxygen binding, respectively. Therefore, there is a well-established biochemical foundation supporting the additive value of 3D protein structure in protein property prediction.

With the recent advances in large pretrained language models (LMs) from natural languages^8–10^, various types of LMs have also been adapted to protein modeling by treating protein sequences as the language of life, where tokens are AAs. Prominent examples include transformer-based LMs such as those developed in ProtTrans^11^ and ESM^12^ as well as long short-term memory (LSTM)-based LMs from Alley et al.^13^ and Heinzinger et al.^14^. Trained with billions of natural protein sequences alone in self-supervised fashion, protein LMs have been shown to achieve state-of-the-art performance on various residue-level and protein-level tasks^11,12^.

Mechanistically, researchers found LMs are able to learn evolutionary information embedded in billions of protein sequences across many species. Concretely, protein LMs can embed proteins from different domains of life (*archaea*, *bacteria*, and *eukarya*)^11^. More interestingly, a recent study found that protein LMs trained on mostly wild-type (WT) sequences with masked LM objective, can be used to quantify mutational effects using the LM likelihood without further training^15^.

Protein LMs can also learn rudimentary structural information without explicit supervision. For instance, the protein-level embedding from LMs can predict protein structure classes encoded in SCOPe (Structural Classification of Proteins—extended)^16^. Residue-level embeddings from LMs have also been shown to be predictive of secondary structure and tertiary contact map^11,12^, even in few-shot learning settings^17^. Rao et al.^17^ demonstrated that transformer-based protein LMs learn to encode residue contacts in their attention maps.

Protein 3D structures have also been explicitly used for both general-purpose protein LMs and property prediction tasks. Bepler and Berger^18^ developed a bidirectional LSTM model with a residue contact prediction objective to incorporate structural information. For protein property prediction tasks, protein 3D structures have been mostly treated as a graph of AA residues, which can then be fed into graph neural networks (GNNs). By representing protein 3D structures as AA graphs based on contact maps built from C-alpha distances, Villegas-Morcillo et al.^19^ found a graph convolutional net (GCN)^20^ underperformed a sequence-only baseline where AA embeddings from protein LMs are used to predict protein functions. Gligorijević et al. introduced DeepFRI^21^, a GCN-based architecture to combine the information from sequence and structure by incorporating AA embeddings from protein LMs as node features. DeepFRI achieved state-of-the-art performance on various protein function prediction tasks.

However, representing 3D structure by building AA graphs from the contact map is a reductionist approach: it only captures inter-residue distances and interactions while disregarding fine-grained details in protein structures such as residue orientations. Studies explicitly incorporating structural information have achieved improved performance on protein structural modeling. For instance, Ingraham et al.^22^ added residue directions and orientations as edge features on the protein graph to improve the generative modeling of protein sequences from this structural representation. By predicting inter-residue orientations in addition to distances, Yang et al.^4^ also improved their protein structure prediction algorithm. More recently, Jing et al.^23^ developed a novel neural module geometric vector perceptrons (GVP) for learning vector-valued and scalar-valued functions over 3D Euclidean space. The output of GVP is equivariant or invariant to rotations and reflections in 3D Euclidean space. When processing protein graphs with rich node and edge features including dihedral angles, backbone directions, inter-residue distances, GVP achieved state-of-the-art performance on protein design and model quality assessment tasks.

Moreover, protein property prediction methods such as DeepFRI^21^ performs the feature extractions for protein sequence and structures in isolation. Therefore, we reason that the potential synergistic predictive values between structure and sequence cannot be fully exploited by those methods.

In this study, we developed a novel end-to-end deep learning framework for protein property prediction by leveraging information from both protein sequences and 3D structures. Our end-to-end neural architecture organically connects a protein LM and a GNN, allowing gradients to back propagate into both the GNN and the LM. Our approach can be considered a novel fine-tuning procedure for protein LMs by injecting inductive bias from 3D structures, which outperformed previous approaches on protein property prediction tasks that either keep the protein LM fixed^21^, or do not incorporate structural information^11,12^. We also demonstrated that the structural fine-tuning of protein LM improves its ability in assessing mutational effects in a zero-shot fashion.

## Results

### LM-GVP, an end-to-end deep learning framework combining protein structure and sequence using protein LM with a GNN prediction head

LM-GVP, a novel ensemble deep learning framework facilitates joint training of protein LM and GNN with desirable geometric properties, capable of processing more detailed representations of protein structures. LM-GVP (Fig. 1) is composed of a protein LM and a GVP network^23^: the protein LM takes protein sequences as input to compute embeddings for individual AAs, which are then concatenated into the node scalar features in the AA graph representation of protein structures. The GVP network is responsible for learning complex structure-function relationships from the AA graphs, with help from the LM. In the training phase, LM-GVP is trained in an end-to-end fashion: the gradients are back-propagated from the GVP network to the transformer blocks of the protein LM (Fig. 1) so that the parameters in the protein LM can update accordingly to produce optimized embeddings for predicting the intrinsic structural and functional properties of proteins. Our architecture can be considered as a novel fine-tuning method for protein LMs capable of injecting inductive bias from protein structures via the GNN prediction head.

**Figure 1.**
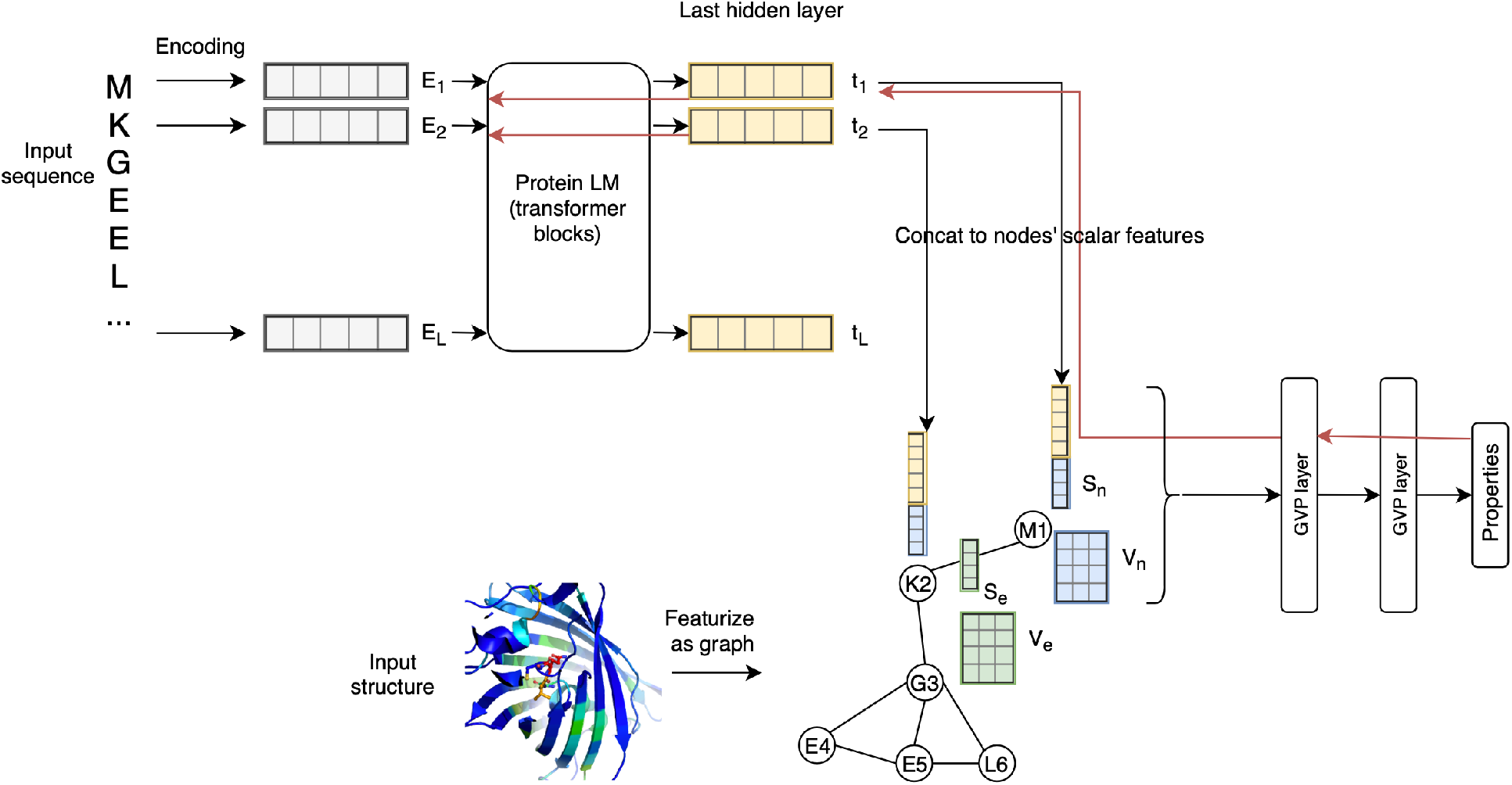
Schematic overview of the LM-GVP, a generalizable deep learning framework for protein property prediction from sequence and structure. LM-GVP is composed of a protein LM connecting the amino acid (AA) embeddings to a GVP network. Protein sequences are processed by the LM to calculate AA embeddings. Protein structures are first featurized to a graph of AAs, with scalar and vector features on both nodes and edges to encode information about distance and direction. The AA embeddings are concatenated with other node features on the graph, which are processed by the GVP network to learn to make predictions about protein properties. The black and red arrows indicate forward and backward passes in the network, respectively.

### LM-GVP improves protein property prediction performance compared to models using protein sequence and structure in isolation

We evaluated the performance of LM-GVP on a collection of 5 publicly available protein property prediction datasets, including three collections of gene ontologies (GO)^24,25^: Cellular Component (CC), Molecular Function (MF), and Biological Process (BP), as well as two protein engineering datasets, Fluorescence and Protease stability from Tasks Assessing Protein Embeddings (TAPE)^26^. To prove the synergistic predictive value between protein sequences and structures, we set up baseline models using sequence-only and structure-only information. The sequence-only baseline fine-tunes the protein LM by connecting a linear layer to the pooled output of the classification token, whereas the structure-only baseline is a GNN network (GVP or GAT) trained on AA graphs with one-hot encoded identity of the AA as scalar node features. We also compared LM-GVP with 2-stage architectures developed in DeepFRI^21^ where protein LM and GNN are trained independently. Overall, we found that the LM-GVP achieved the best performances on all three subsets of the GO dataset (Table 1) and competitive results on Fluorescence and Protease (Table 2) over baseline models and 2-stage models. Consistent with DeepFRI^21^, we found that the 2-stage models combining sequence and structural information outperform sequence-only and structure-only baselines across all three GO subsets (Table 1). In addition, we demonstrated that the GVP network outperforms GAT in both structure-only baselines and 2-stage models, suggesting the rich node and edge features in AA graphs are informative to protein properties and that the GVP network is able to leverage that information. The observation that our LM-GVP architecture, when trained in 2-stage procedure, underperforms the ones trained end-to-end (Table 1, 2), indicates that back-propagating the gradients to the protein LM indeed boosts the performance of various protein property prediction tasks.

**Table 1.** Performance of LM-GVP and other baseline models in predicting protein functions in GO hierarches. Function-centric and AUPR and protein-centric Fmax scores from hold-out test sets are shown in the table to evaluate the predictive performances. GAT: graph attention network; GVP: geometric vector perceptron network.

| Method category | Method | GO-CC |  | GO-BP |  | GO-MF |  |
| --- | --- | --- | --- | --- | --- | --- | --- |
|  |  | AUPR | F<sub>max</sub> | AUPR | F<sub>max</sub> | AUPR | F<sub>max</sub> |
| **Sequence-only** | ProtBERT fine-tune | 0.234 | 0.408 | 0.188 | 0.279 | 0.464 | 0.456 |
| **Structure-only** | GAT | 0.249 | 0.385 | 0.171 | 0.284 | 0.329 | 0.317 |
|  | GVP | 0.278 | 0.420 | 0.224 | 0.326 | 0.458 | 0.426 |
| **Sequence+structure** | LM-GAT (2-stage) | 0.353 | 0.494 | 0.277 | 0.393 | 0.529 | 0.507 |
|  | LM-GVP (2-stage) | 0.413 | 0.515 | 0.278 | 0.386 | 0.544 | 0.507 |
|  | LM-GVP | **0.423** | **0.527** | **0.302** | **0.417** | **0.580** | **0.545** |

**Table 2.**
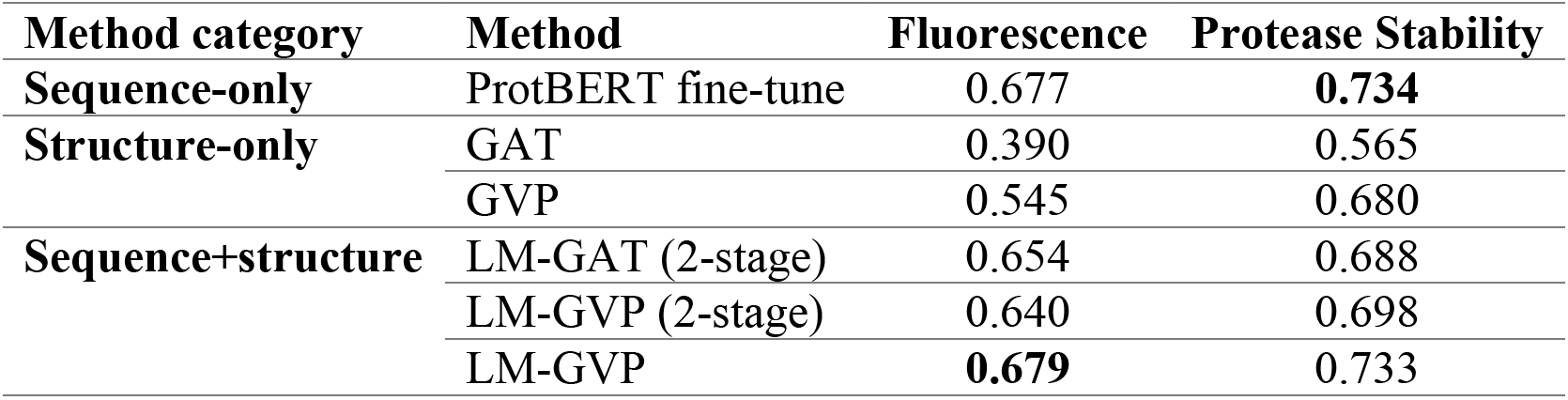
Performance of LM-GVP and other baseline models in TAPE protein engineering tasks. Spearman’s correlation coefficients calculated from hold-out test sets are shown in the table to evaluate the predictive performance of the regression tasks.

| Method category | Method | Fluorescence | Protease Stability |
| --- | --- | --- | --- |
| **Sequence-only** | ProtBERT fine-tune | 0.677 | **0.734** |
| **Structure-only** | GAT | 0.390 | 0.565 |
|  | GVP | 0.545 | 0.680 |
| **Sequence+structure** | LM-GAT (2-stage) | 0.654 | 0.688 |
|  | LM-GVP (2-stage) | 0.640 | 0.698 |
|  | LM-GVP | **0.679** | 0.733 |

Next, we examined the predictive performances of GO terms by different methods in order to delineate the predictive power of LM-GVP on different protein functions (Fig. 2). We found that GO-MF terms are more predictable (micro-AUPR = 0.580) than GO-CC (micro-AUPR = 0.423) followed by GO-BP (micro-AUPR = 0.302) (Table 1 and Fig. 2A). The trend is consistent across all predictive models (Table 1 and Fig. 2A). The vast majority of GO terms (2263 out of 2737, or 82.7%) can be better predicted by the LM-GVP than baselines. When attributing the predictability of protein functions to sequence and structural information, we discovered that enzymatic activities such as lysozyme activity (GO:0003796), DNA topoisomerase II activity (GO:0003918), and racemase/epimerase activity (GO:0016855), are much more predictable by protein sequences than structures (Fig. 2B, Table S1). This corresponds to structural plasticity, but sequence invariance often found in enzyme functional sites^27,28^. Protein functions that are more predictable by structures than sequences are enriched for components of large protein complexes such as viral envelope (GO:0019031), photosystem I reaction center (GO:0009538), hemoglobin complex (GO: 0005833), and MHC protein complex (GO:0042611) (Table S2). Components of large protein complexes typically have distinctive structural motifs that are easily enhanced with the LM-GVP model. We also identified GO terms with lower predictability than sequence-only or structure-only baselines (Fig 2C-D, Table S3–4). Furthermore, we noticed sequence and structure information combined does not necessarily outperform every individual property prediction task including Protease stability (Table 2). The predictive performance of the sequence-only model is also competitive with LM-GVP (Table 2).

**Figure 2.**
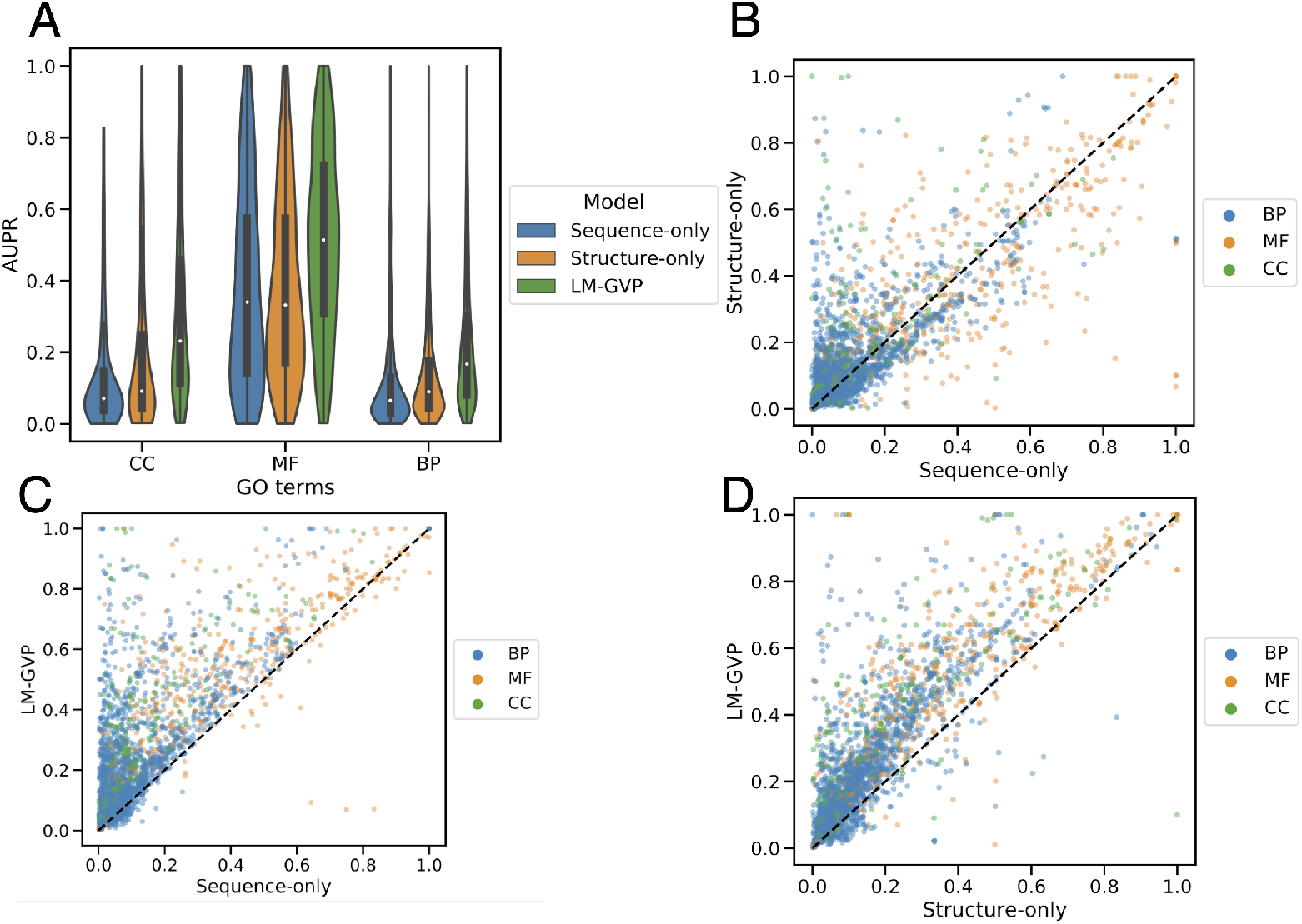
Distribution of model performance across individual GO terms. (A) Violin plots showing the distribution of Area Under the Precision Recall curve (AUPR) over subsets of GO terms predicted by different models. (B-D) Scatter plots of AUPR values across GO terms from different models. Each dot in the plots corresponds to an individual GO term, colored by its subset indicated in the legend. BP: biological process; MF: molecular function; CC: cellular component.

### Integrated gradient provides residue-level interpretability for LM-GVP

Protein functions are often exhibited by specific regions on the 3D protein structures, such as active sites/regions of enzymes and binding interface between ligands and receptors. We next examined whether LM-GVP can attribute the positive predictions of certain protein functions to the active residues known to be mechanistically essential. We applied Integrated Gradient (IG)^29^, a model-agnostic method for interpreting neural models. For each protein, IG generates a saliency map where each residue is associated with an attribution score, indicating how much the residue contributes to the model’s prediction. We calculated AUROC to quantify the agreement between the saliency scores and binding sites. We found that LM-GVP can identify the active sites on proteins responsible for the binding ATP, GTP and Heme (Table 3, Fig. S1) more accurately than sequence-only and structure-only baselines. The relatively lower AUROC for 4ZLT-F and 3WCY-I (which are protein molecules that bind to cytokine receptors) can be explained by the nature of their large, relatively less well-defined, and flexible cytokine-binding surfaces (Fig. S1). This is more challenging for LM-GVP’s IG-derived saliency scores to perfectly capture.

**Table 3.** Area under the receiver operating characteristic curve (AUROC) quantifying the agreement between saliency scores and known active sites responsible for respective molecular functions (MF). (x) indicates a false prediction.

| **Protein** | **GO Molecular Function** | **ProtBERT fine-tune** | **GVP** | **LM-GVP** |
| --- | --- | --- | --- | --- |
| **1E2Q-A** | ATP binding | 0.718 | 0.570 | **0.780** |
| **1Z83-A** | ATP binding | 0.543 | 0.496 | **0.670** |
| **6IF2-B** | GTP binding | 0.550 | 0.515 | **0.638** |
| **2GF0-A** | GTP binding | 0.470 | 0.420 | **0.541** |
| **3HF4-B** | heme binding | 0.412 | 0.440 | **0.616** |
| **3IBD-A** | heme binding | 0.428 | 0.402 | **0.578** |
| **4ZLT-F** | cytokine receptor binding | 0.633 (x) | 0.518 | **0.516** |
| **3WCY-I** | cytokine receptor binding | 0.448 (x) | 0.478 (x) | **0.545** |

We further extracted the latent representations of the proteins at the penultimate layer of LM-GVP and performed clustering analysis. The latent representations are first projected onto a 2D plane using UMap^30^, followed by DBSCAN^31^ to detect and visualize families of proteins (Fig. 3A). By inspecting proteins within these clusters, we demonstrated that LM-GVP’s latent representation is able to identify both sequence and structural features known to be important for ATP binding. For instance, the saliency scores highlight a cluster of ATP-binding proteins with Walker A motif (GxxGxGK[S/T]) (Fig. 3B). Interestingly, we found an apparent outlier, 2ORV-A human thymidine kinase 1 (*TK1*), within this cluster on the MSA with distinct sequence and an additional high saliency region at positions 315-320 (Fig. 3B). When aligning 2ORV-A with a typical member of this cluster, 1E2Q-A human thymidylate kinase encoded by *DTYMK*, we found both their structures and salient regions are well aligned (Fig. 3C). Similar structural alignments are also observed with other proteins in this cluster (Fig. 3D). This result suggests that LM-GVP, although not explicitly trained to perform residue-level tasks, is able to learn from both structural and sequence features associated with functions to make accurate predictions.

**Figure 3.**
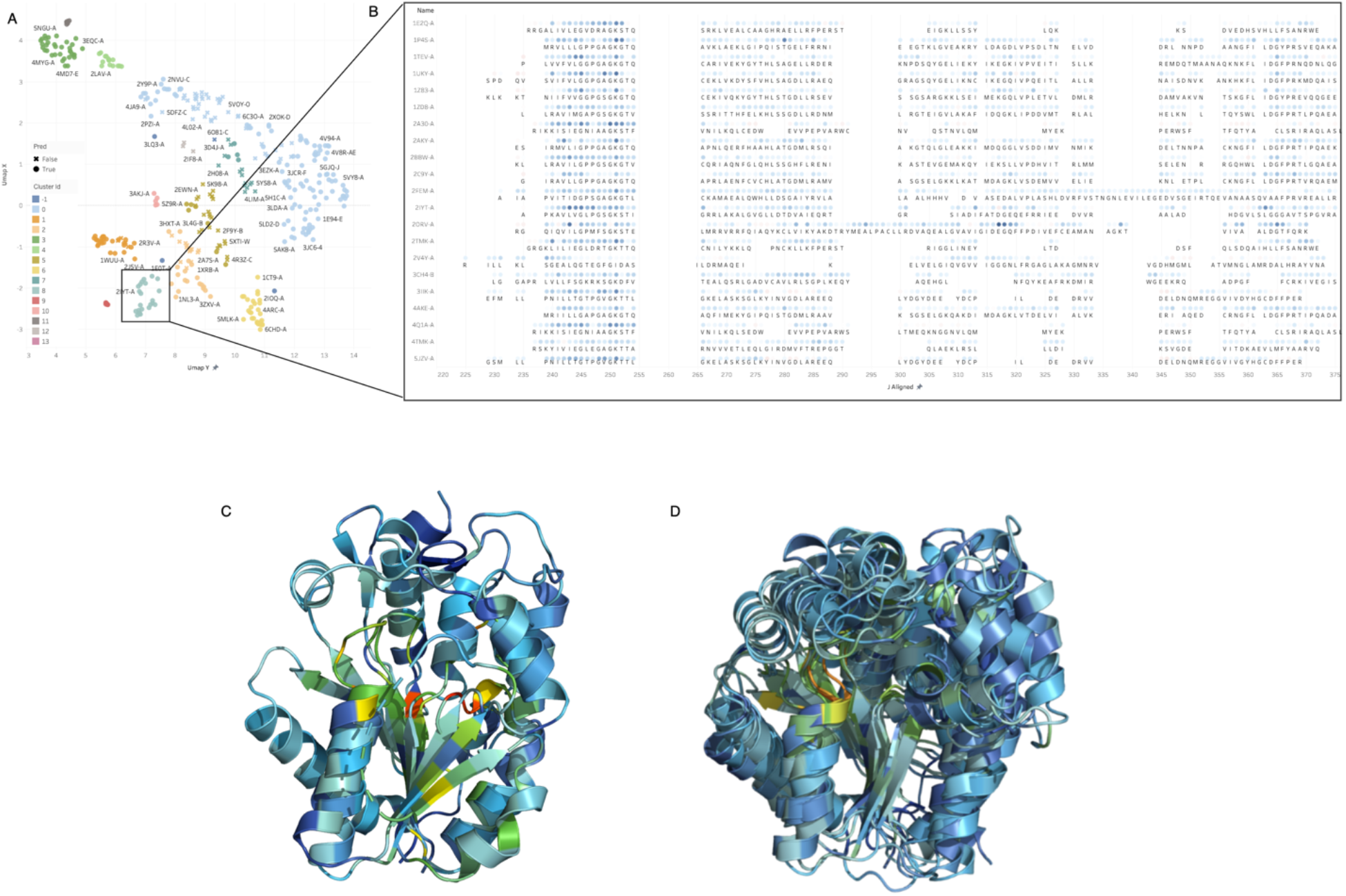
Interpretation of LM-GVP on predicting ATP binding proteins. (A) UMAP projection of the LM-GVP representation shows the clusters in ATP binding proteins. (B) A selected cluster of proteins and their saliency maps aligned using Multiple Sequence Alignment (MSA). The visualization shows common active sites/regions responsible for ATP binding. (C) Structural alignment between 2ORV-A (human thymidine kinase 1) and 1E2Q-A (human thymidylate kinase) with residues colored by saliency scores. (D) Structural alignment of proteins in (B), with residues colored by saliency scores.

### Exploring the effects of fine-tuning protein LMs with GVP

LM-GVP can be regarded as a novel fine-tuning procedure for protein LMs: it explicitly injects the inductive bias from complex structure-function relationships into the protein LM. We next seek to gain more mechanistic insights into LM-GVP by exploring the effects of our novel fine-tuning approach in protein LMs with regards to two important tasks. Specifically, we asked if the structural inductive bias helps improve the expressiveness of the protein LMs in assessing mutational effects and predicting contacts.

### Fine-tuning protein LMs with protein structures enhance their zero-shot prediction for mutational effects

We perform zero-shot analysis of our models to assess the degree to which the transformer component of LM-GVP internalizes information about the relative likelihood of each amino acid in the context of the protein’s overall sequence. The first row of Table 4 provides the value of Spearman’s rank correlation coefficient (rho) for the ProtBERT language model out of the box (i.e., not fine-tuned on any particular downstream task). We obtain values for 6 assays across deep mutational screens for 4 different proteins and observe that ProtBERT’s internal representation of the contextualized amino acid distributions is greater than 0.5 for PABP in Yeast as well as log fluorescence of GFP in *Aequorea victoria*, whereas these distributions are relatively uninformative in the case of the K_m_ value of BLAT in *E. coli* (rho=0.05). We replicate this analysis for six additional models: for each of the three available GO classification tasks, we fine-tune ProtBERT directly in addition to fine-tuning LM-GVP. As expected, most of the transformer models fine-tuned directly on the GO data subsequently produce lower values of rho for a given mutational screen than ProtBERT, whereas the likelihoods produced by the LM-GVP models generally retain or exceed their correlation with the target values. This is the expected result of relative over-fitting of the transformer weights to the specific GO task during fine-tuning with a simple linear prediction head. In contrast, the geometrically-aware prediction head of LM-GVP is able to leverage the protein structure associated with a given sequence to obtain better classification performance on each task while simultaneously preserving important information about the amino acid distributions underlying the language model. This is likely the direct result of the fact that the protein’s tertiary structure is the mediating factor through which all proteins ultimately perform their function. Critically, we see that the transformer fine-tuned directly on the GO-MF labels perform worse than their out-of-the-box counterparts in terms of the rank correlation on all 6 zero-shot analyses (Table 4), while the LM-GVP model fine-tuned on this same dataset performed better for all 6 (Table 4). This validates the hypothesis that the GVP head of the model is particularly valuable when supervising the model with information critical to protein function and therefore heavily dependent on structure.

**Table 4.** Zero-shot performance of protein LMs in predicting mutational effects. Spearman’s correlation coefficients are shown in the table across datasets and protein LMs underwent different fine-tuning methods and tasks.

| Fine-tune | Fine-tune targets/Datasets | PABP Yeast (log) | DLG4 Rat (CRIPT) | DLG4 Rat (Tm2F) | BLAT *E. coli* (Km) | BLAT *E. coli* (Vmax) | GFP (log fluorescence) |
| --- | --- | --- | --- | --- | --- | --- | --- |
| None | N/A | 0.574 | 0.456 | 0.242 | 0.057 | 0.249 | 0.613 |
| Seq-only | GO-BP | 0.082 | 0.064 | 0.025 | 0.026 | 0.266 | 0.108 |
|  | GO-MF | 0.040 | 0.193 | 0.163 | 0.053 | 0.244 | 0.202 |
|  | GO-CC | 0.276 | 0.203 | 0.185 | 0.037 | **0.361** | 0.415 |
| LM-GVP | GO-BP | 0.554 | 0.438 | 0.220 | **0.068** | 0.335 | **0.627** |
|  | GO-MF | **0.585** | 0.466 | 0.245 | 0.062 | 0.290 | 0.620 |
|  | GO-CC | 0.569 | **0.467** | **0.252** | 0.064 | 0.300 | 0.616 |

### LM-GVP helps preserve the structural information in protein LMs

Many studies^12,17,32^ have identified the connection between attention maps in protein LMs and proteins’ structural features. Rao et al.^17^ further illustrated that the attention maps of transformer-based protein LMs trained on billions of sequences with the unsupervised LM objective are able to learn protein contact maps in few-shot settings, suggesting some structural information is encoded in the pretrained protein LMs, even without explicitly providing it during training. Since LM-GVP explicitly combines structural information with protein LMs, we next explored how the intrinsic structural information is changing over the course of LM-GVP fine-tuning and sequence-only fine-tuning of protein LMs for different property prediction tasks.

To assess the amount of structural information represented in our protein LMs, we followed Rao et al.^17^ to first learn a L1-logistic regression model to predict proteins’ contact maps using the self-attention maps from the LM (Fig. 4A). Consistent with Rao et al.^17^ findings, the transformer heads that resemble the contact maps are mostly concentrated on the last layers of ProtBERT (Fig. 4B). We next quantified the precision for predicting contacts using attention maps from ProtBERT with and without fine-tuning over 5 tasks. ProtBERT without fine-tuning for any property prediction tasks can predict the contacts in the GFP proteins significantly better than proteins from the GO dataset (Fig. 4C), suggesting that ProtBERT already has better knowledge about the structures of GFP proteins compared to the diverse collection of proteins in the GO dataset. This observation further suggests that the additive predictive value from protein structures might not be as prominent, if any, on the Fluorescence and Protease datasets compared to the GO dataset, as shown by our experiments (Table 1, 2). Interestingly, we also found after fine-tuning with protein sequences alone on all 5 tasks, the protein LM sacrifices the intrinsic knowledge it stored about protein contact maps to improve predictive performance on the property prediction (Fig. 4C). This effect is more pronounced on specialized tasks such as predicting fluorescence. In contrast, protein LM after LM-GVP fine-tuning maintains its representation power of protein contact maps among three GO tasks significantly better than fine-tuning with sequence alone (Wilcoxon signed-rank test p-values = 5.31e-5; 3.15e-27; 2.01e-34, in CC, BP, MF tasks, respectively). However, LM-GVP fine-tuned LMs are not significantly better at preserving the contact map information on the protein engineering tasks. This result indicates that LM-GVP are generally better at preserving the structural representations within protein LMs while optimizing the performance at predicting protein properties. The loss of representation power in LM also gives us a hint on the performance of LM-GVP for different tasks. However, it is worth noting that residue contact map is merely one aspect of information contained in protein 3D structures.

**Figure 4.**
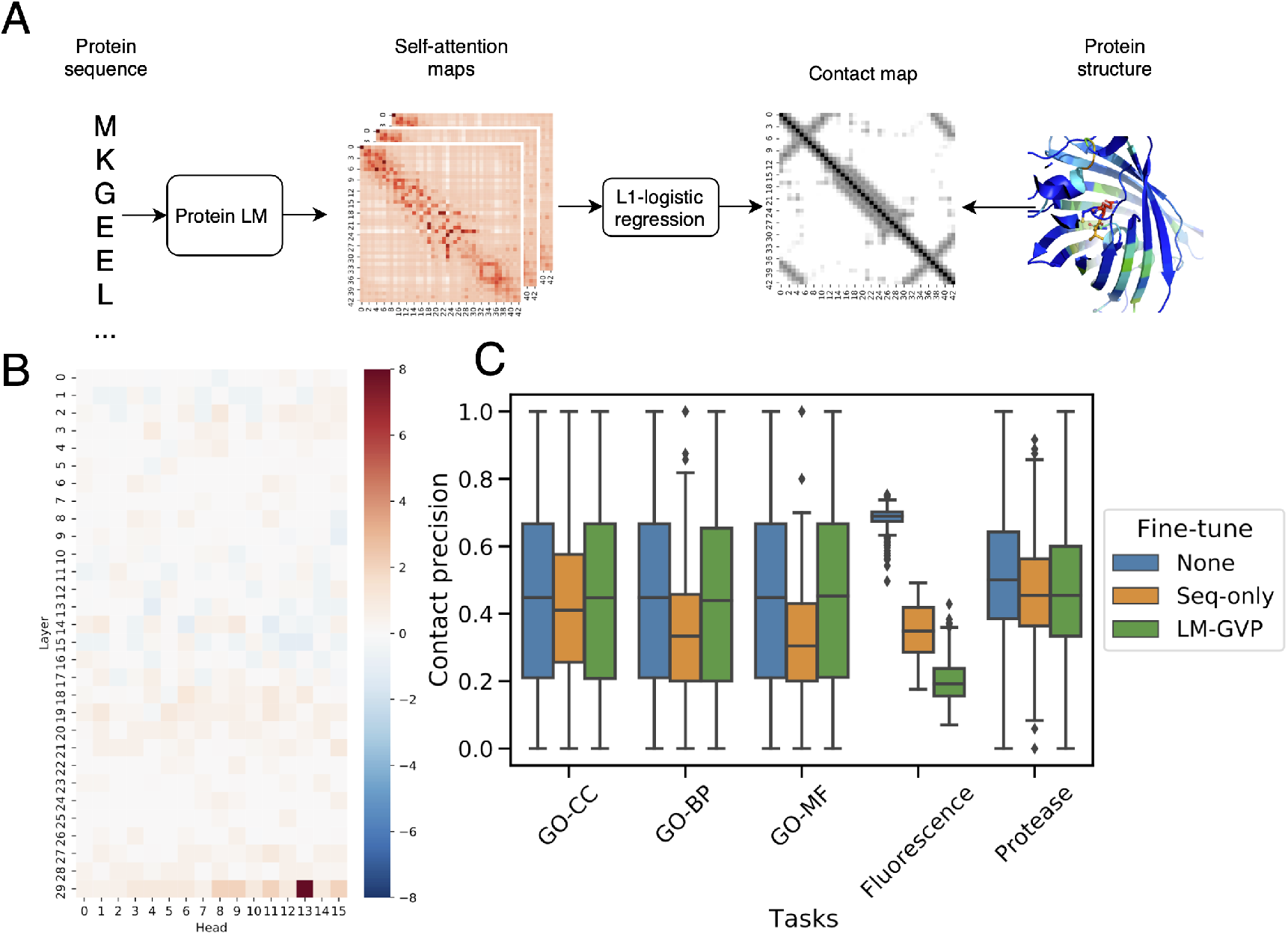
Predicting protein contact maps with self-attention maps from protein LMs. (A) Schematic view showing how self-attention maps from protein LMs can be used to predict contact maps. (B) Weights from L1-logistic regression trained to predict residue contacts. (C) Box plot showing the distribution of contact map prediction precision from self-attention maps in the protein LM before and after different fine-tuning methods.

## Discussion

The 3D structure ultimately dictates a protein’s function: if the structure of a protein is altered, so is its function (e.g., mis-folding). However, due to the limited availability of high-resolution 3D structures, protein LMs trained solely on 1D sequence information dominate many predictive applications related to proteins.

In this study, we demonstrated the additive value of incorporating structural information on top of protein LMs for property prediction tasks. Our LM-GVP leverages fine-grained structural information beyond just binary contact maps, providing a novel inductive bias to instruct the pretrained protein LM to jointly learn with its GVP head to optimize the predictive performance for protein properties. Our observation that structural information is complementary to protein LMs also suggests that the representation capacity of protein LMs for structure is limited. After all, most protein LMs are trained on protein sequences alone and the objective functions (e.g., masked LM objective and auto-regressive objective) may not encourage the LMs to learn complex evolutionary sequence-structure relationships. One potential solution to increase the structural representation of protein LMs is to jointly learn from sequence and structure as pioneered by Bepler and Berger^18^, where contact maps were explicitly used when training the LM. Graph transformers^33^ could also be leveraged to learn from rich attributed graphs constructed from 3D structures in self-supervised settings. However, researchers still need to tackle the challenge of relatively fewer available protein structures (∼180K) compared to sequences (>300B).

LM-GVP can be also considered a novel fine-tuning procedure for protein LMs vis-à-vis building upon the notion that LMs capture “the language of life”. Such a procedure can be extrapolated to natural languages as well. Natural language can also be represented as graphs in various ways, such as dependency graphs (e.g. syntactic dependency parsing tree or semantic dependency parsing tree) and co-occurrence graphs^34^. On text classification tasks, LM-GVP can be adopted to back propagate the gradients from a GNN operating on graphs of tokens to LMs to potentially improve predictive performance. Other applications of natural language processing (NLP) such as question-answering have also seen a similar approaches^35^ that organically combine GNNs with LMs.

In conclusion, we described the novel method of LM-GVP, a generalizable deep learning framework for protein property prediction, harnessing the representation power from pretrained protein LMs and fine-grained structural information to achieve the state-of-the-art performance on various property prediction tasks. We have also provided mechanistic insights into the impact of structurally-instructed fine-tuning for protein LMs and believe there could be beneficial implications in natural languages.

## Methods

### Machine learning models for protein property prediction

We experiment with many machine learning models for protein property prediction tasks, including sequence-only and structure-only baselines, 2-stage methods for combining of sequence and structural information, as well as our LM-GVP. Here we describe those models in great details.

#### Sequence-only baseline

protein property prediction is analogous to document classification tasks in natural language. To use protein LMs for property prediction, we fine-tune ProtBERT^11^ with a dense layer with number of output units corresponding to different tasks. The dense layer is connected to the classification token [CLS] of ProtBERT, which is the last layer hidden-state of [CLS] further processed by a linear layer and a Tanh activation function. During fine-tuning, the gradients are back-propagated the to all the layers in ProtBERT except for the embedding layer.

#### Structure-only baselines

Graph neural networks (GNNs) have been used for protein property predictions^19,21^. Here we describe our structure-only baselines based on two types of GNNs: GAT and GVP.

For both GAT and GVP, the 3D structure of protein is transformed to a proximity graph 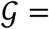 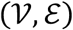 following Ingraham et al.^22^ and Jing et al.^23^. Each node 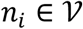 in the proximity graph 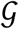 corresponds to an amino acid (AA) and has node features 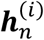, where *i* ∈ 1,2, …, *L* denotes the indices of AA in the protein sequence of length *L*. Edges in the graph 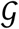 connect adjacent AAs and has edge features 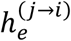 Edges are formed by connecting k-nearest neighbors for each node based on the Euclidean distance from C-alpha coordinates ***X*** ∈ ℝ^*L*×3^ with *k* = 30 for all experiments.

Node features 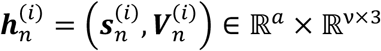 are composed of scalar and vector features. Scalar features 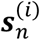 include the sines and cosines of the dihedral angles ϕ, Ψ, ω; the one-hot representation of AA identity. Vector features 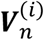 are consist of the forward and reverse unit vectors in the directions of adjacent C-alpha atoms from two neighboring AAs; the unit vector in the imputed direction of C-alpha and C-beta atoms.

Edge features 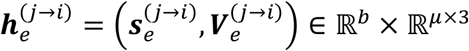 are composed of scalar and vector features as well. Scalar features 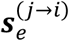 include the encoding of C-alpha distance in terms of 16 Gaussian radial basis functions with centers evenly spaced between 0 and 20 angstroms; a positional encoding of *j* − *i* as described in Vaswani et al.^8^, representing the AA distance alone the 1D protein sequence. The unit vector in the direction of connecting C-alpha atoms is used as vector features 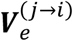.

Both GAT and GVP follow the message passing paradigm^36^ where messages are computed from neighboring nodes and edges, which are subsequently used to update node embeddings at each graph propagation step. Generically, a message on a given edge *j* → *i* is first computed by:

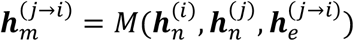

Next, the node embedding is updated by aggregating the messages from all of its edges:

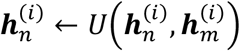

where 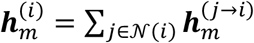 sums up the messages, and 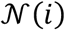 denotes the neighbors of n_*i*_ in the graph 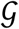.

GAT and GVP differ by the choice of message function *M* and update function *U*, as well as their respective inputs. To reproduce the settings from DeepFRI^21^, the GAT network only uses the one-hot encoding of the AA’s identity as node features and its message function is defined by:

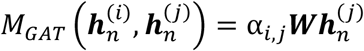

where α_*i*,*j*_ is the learned attention weight. Its update function is defined by:

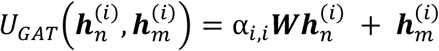

3 layers of GAT convolutions are used on the protein graphs with different number of AAs. To aggregate the node embeddings to the final protein-level representation, we first concatenate node features from all layers into a single feature matrix, i.e., ***H*** = [***H***^(1)^, ***H***^(2)^, ***H***^(3)^] ∈ *R*^*L*×*C*^, and then perform a global sum pooling layer over the AA axis to obtain a fixed vector representation.

The GVP network is modified from the model quality assessment (MQA) network described by Jing et al.^23^. It is composed of three consecutive GVP convolution layers, which operates on the tuple of scaler and vector features:

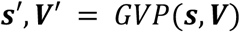

The GVP network also take advantage of the both node and edge features described above when computing the message:

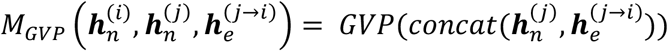

The update function of GVP convolution layer is defined by:

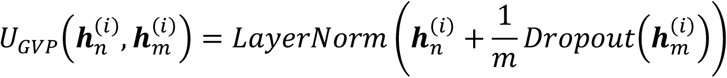

where *m* is the number of incoming messages. In the readout phase, similar to the GAT network, the GVP network also concatenate the node representations from the three layers, separately for scalar and vector embeddings, and then use a global average pooling layer over the AA axis to obtain a fixed vector representation.

#### 2-stage models

we implement 2-stage models following Gligorijević et al.^21^ to combine information from protein sequence and structure using AA embeddings from protein LM and GNNs, respectively. This is a 2-stage method works by calculating the AA embeddings from the protein LM in the first stage:

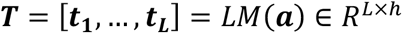

where ***a*** = [***a***_**1**_, …, ***a***_***L***_] is the sequence of AA tokens and *h* denotes the hidden layer dimension in the LM. Next, the AA embeddings ***t***_*i*_ are used as node scalar features for the GNNs in the second stage for learning protein-level properties, replacing the one-hot encoding described previously. The resultant node features become

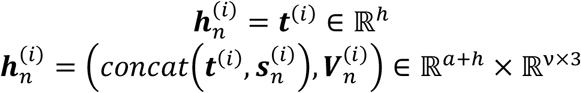

for GAT and GVP networks, respectively. The GNNs used in the 2-stage model are identical to those used in structure-only baseline described above. We use pretrained ProtBERT developed in^11^ as the protein LM across all experiments.

#### LM-GVP

is our novel method with the same architecture as the 2-stage model where GVP network is used as the GNN module, but trained in a 1-stage end-to-end fashion. That is, the gradients are back-propagated into the protein LM’s transformer layers. In the training phase, we adopted the gradual unfreezing technique developed by Howard and Ruder^37^ by first learn the parameters in the GVP network while keeping the parameters in the LM frozen until converge. Then we unfreeze the parameters in the LM’s transformer layers to fine-tune the parameters in both the LM and the GVP network.

#### Details on model training

We use mean squared error (MSE) and weighted cross-entropy loss function for regression and multi-label classification tasks, respectively. To account for class imbalance in multi-label classification settings, we weight the binary cross-entropy losses from each label *j* based on the inverse of the positive instance frequency:

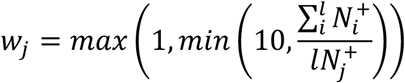

where 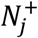 denotes the number of positive instances for label *j* and *l* denotes the number of labels.

Key hyperparameters including learning rate and batch size across experiments are determined through grid search in the choices of [1e-4, 1e-5, 1e-6] for learning rate and batch size of [16, 32] based on model’s loss function on validation set. To avoid overfitting, we employ an early stopping criterion with patience = 10 epochs and trained for a maximum of 200 epochs. ADAM optimizer^38^ with β_1_ = 0.9, β_2_ = 0.999 is used for optimizing the learnable parameters. All models are implemented using the Pytorch deep learning library and training are performed using Pytorch-Lightning library with 16-bit mixed precision training using 8 NVIDIA V100 GPUs with 16 or 32 GB of memory each on Amazon SageMaker.

### Datasets and tasks for protein property prediction

We obtain 3 datasets covering different tasks for protein property prediction. The Gene Ontology (GO) dataset contains three hierarchies of protein functions: biological processes (BP), molecular functions (MF), and cellular components (CC). This dataset is the downloaded from https://github.com/flatironinstitute/DeepFRI/tree/master/preprocessing/data provided in DeepFRI^21^. The construction, preparation, and train/valid/test splitting strategy for of this dataset are comprehensively described in DeepFRI^21^. We downloaded the protein structures in PDB format for proteins in the GO dataset from RCSB PDB^39^ and transform the protein structure to attributed graphs as described in a previous section (Structure-only baselines). The three tasks within the GO dataset are multi-label classification.

We also obtain two protein engineering datasets, Fluorescence and Protease stability from TAPE^**26**^. The Fluorescence dataset contains green fluorescent protein (GFP) with mutations and corresponding log-fluorescence intensity. The goal of this task is to predict the log-fluorescence intensity from the sequence and/or structure of the GFP mutants. We adopt the same train/valid/test splitting strategy from TAPE^**26**^. The Protease stability dataset contains proteins and measurement of their intrinsic stability in terms of maintaining its fold above a protease concentration threshold. We split the train/valid/test sets for the Protease stability dataset by stratifying the regression target. Both the Fluorescence and Protease stability datasets are regression tasks with single target. The 3D structures of proteins in the Fluorescence and Protease datasets are generated by Rosetta first by introducing the mutations with the fixbb protocol^**40**^ and then relaxing^**41**^ the resulting structure.

### Model interpretation

Integrated Gradients (IG)^29^ is a model-agnostic interpretation technique that attributes the prediction of a model to its input features. It can be applied to any differentiable model and does not require modification of the model structure. The method has been widely used in interpreting DNNs for natural language understanding and motivated by its success, we also apply it here to obtain residue-level importance for predicting molecular functions.

IG integrates the gradient along a straight-line path between a baseline input and the original input to obtain the feature attributions / saliency scores. More specifically, if we denote the original input as ***x***, the baseline input as ***x***′, and the model under analysis as ***F***, IG along the ***i***^***th***^ dimension of the input can be calculated as follows:

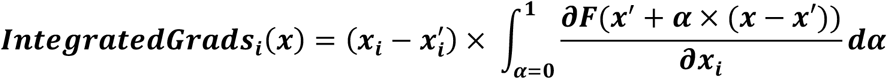

To analyze the LM-GVP model, we used the embedding of a sequence that consists of entirely neutral tokens ([SEP]) as the baseline input. We denote the original AA sequence as ***α***′ and the modified sequence with only [SEP] tokens as ***α***′. Their embeddings are calculated respectively as ***x*** = ***LM***(***α***) ∈ ***R***^***L***×***h***^ and ***x***′ = ***LM***(***α***′) ∈ ***R***^***L***×***h***^. The IG attribution can be obtained for each residue in the sequence on each embedding dimension, which we denote as ***ig***(***i***, ***j***), where ***i*** ∈ {**1**, …, ***L***} and ***j*** ∈ {**1**, …, ***h***}. The final saliency score for the ***i***th residue can be calculated by summing over the embedding dimension: 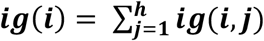 Same approach is used for interpreting the sequence-only baseline. For the structure-only baseline, we also use the residue token embeddings to perform the analysis.

We analyzed the resulted saliency score by comparing it with known binding sites retrieved from BioLiP^**42**^ (https://zhanglab.dcmb.med.umich.edu/BioLiP/download.html). Where data was not available, a sphere of 5.0 was created around the ligand in the binding pocket and all sidechain interactions with the ligand were included. We use the known binding sites to obtain a binary profile for each protein. Each residue is associated with a 0 or 1 ground truth label indicating whether it is known binding sites. Our hypothesis is that if a residue has a higher saliency score, it is more likely to be a binding site. We compute the area under the ROC curve (AUROC) to analyze the alignment between the saliency score obtained via IG and the ground truth binding sites for different molecular functions include ATP binding, GTP binding, heme binding and cytokine receptor binding. See Supplementary Table S5 for results on more proteins.

Uniform Manifold Approximation and Projection (UMAP)^**30**^ is a non-linear dimension reduction technique that can be used to visualize the clusters within high-dimensional data. We applied UMAP to analyze the latent representation at the penultimate layer of LM-GVP (with 400 dimension) and identified families of proteins with similar structural / sequence motifs that are related to their molecular functions (GO-MF terms). We use DBSCAN^**31**^ to extract and analyze selected protein clusters in more detail. Fig 3A shows the dimension reduction and clustering results for a set of proteins with ATP binding function. We select a small cluster for detailed analysis by display the saliency maps of the proteins in the cluster. To facilitate pattern identification, the sequences are aligned using MSA implemented in Biopython^**43**^.

We obtained each protein’s 3D structure from the Protein Data Bank^**39**^ and load it into PyMOL^**44**^. We then used the spectrum coloring command in PyMOL to assign color to each of the residues based on their saliency scores with the most salient residues colored in shades of red and the least salient residues in shades of blue.

### Analysis of mutational effect

We analyze zero-shot performance of our fine-tuned language models on four separate datasets of mutational scans for individual proteins, three of which were provided as part of the DeepSequence GitHub repository^45^ and the fourth as part of TAPE^26^. For each non-wildtype protein sequence, we mask the mutated amino acid(s) and pass the tokenized representation through the transformer component of LM-GVP to generate probability distributions over all possible amino acids at the masked positions. As in Meier et al.^15^, we then compute the masked marginal probability score as

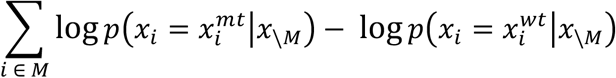

which is the sum of the differences in log-probability of the mutant and wildtype amino acids over all mutated positions in the sequence. We finally calculate Spearman’s rank correlation coefficient between these scores and the original assay values.

### Contact-map prediction analysis

To assess the structural information intrinsic to protein LMs, we adopt the few-shot learning approach described in Rao et al^17^. Briefly, we first calculate the self-attention maps from the pretrained ProtBERT without any fine-tuning on 20 randomly selected proteins with more than 30 AAs, to predict residue contacts defined by C-alpha distance <= 10 Å between residues at least 6 AAs apart to ignore local contacts. The self-attention maps from all layers of the transformer heads were used as features to learn a logistic regression model with L1 penalty to predict contacts. We then fit parameters in the logistic regression model via scikit-learn^46^. In the inference phase, we predict the contact maps for 500 randomly sampled proteins in the test sets of the five datasets. Then compute the precision score between the contact maps predicted from attention maps and the ground truth.

## Acknowledgements

We thank George Karypis and Zheng Zhang from AWS for insightful conversation on graph neural network algorithms.

## Author contributions

Z.W., S.A.C, R.B., P.X., S.P.P., and P.M.C. wrote the manuscript with input from the all authors. S.P.P. and P.M.C. supervised the research. Z.W., S.A.C. and R.B. led the research. S.A.C., R.B., M.R.C, G.P. C.J.W., and P.M.C. conceived the protein property prediction project. Z.W. led the LM-GVP design. Z.W., R.B., M.R.C., and P.X. performed the training and evaluation of neural models. S.A.C. prepared the structure datasets. P.X. led the model interpretation with the input from S.A.C., E.O.S. and N.G.. R.B. performed the zero-shot analyses. M.R.C. and Z.W. performed the contact map analyses. G.P. provided compute infrastructure support. All authors reviewed the manuscript and approved it for submission.

## Data availability

Datasets used in this study are available to download at: https://github.com/flatironinstitute/DeepFRI/tree/master/preprocessing/data and https://github.com/songlab-cal/tape.

## Code availability

The source code for training end-to-end models, together with the neural network weights are available for research and non-commercial use at https://github.com/aws-samples/lm-gvp.

## Supplementary Information

**Figure S1.**
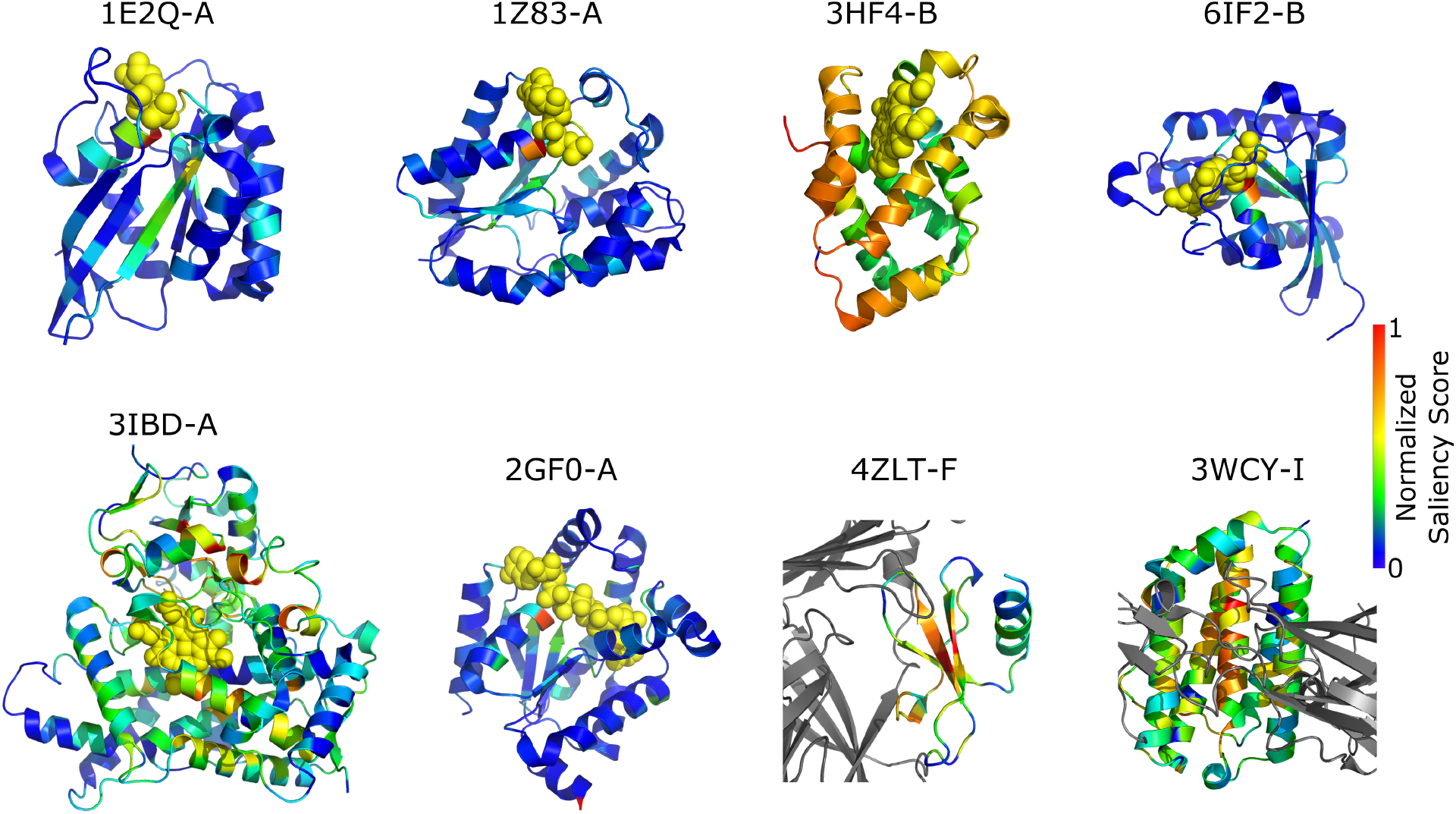
Identification of catalytic residues in enzymes based on saliency scores. For the first six complexes (i.e., 1E2Q-A to 2GF0-A), ligands are shown in yellow spheres while the residues of the receptors they bind to are colored based on saliency scores. In 4ZLT-F and 3WCY-I, the receptors are shown in grey and the residues of the protein ligands are colored based on saliency scores. All the saliency-score-based coloring are done such that the most salient residues are in shades of red and the least salient in shades of blue.

**Table S1.** AUPRs of GO terms with better predictability from sequence-only over structure-only model.

| GO | GO term | task | Sequence-only | Structure-only | Sequence+Structure |
| --- | --- | --- | --- | --- | --- |
| GO:0003707 | steroid hormone receptor activity | MF | 1 | 0.066666667 | 1 |
| GO:0004308 | exo-alpha-sialidase activity | MF | 1 | 0.1 | 1 |
| GO:0016997 | alpha-sialidase activity | MF | 1 | 0.1 | 1 |
| GO:0016855 | racemase and epimerase activity, acting on amino acids and derivatives | MF | 0.833333333 | 0.152631579 | 0.071794872 |
| GO:0036361 | racemase activity, acting on amino acids and derivatives | MF | 0.75 | 0.233333333 | 0.069411765 |
| GO:0003777 | microtubule motor activity | MF | 0.5034053 | 0.002587933 | 0.505068241 |
| GO:0003916 | DNA topoisomerase activity | MF | 1 | 0.5 | 1 |
| GO:0003918 | DNA topoisomerase type II (double strand cut, ATP-hydrolyzing) activity | MF | 1 | 0.5 | 1 |
| GO:0003796 | lysozyme activity | MF | 1 | 0.505256648 | 0.854166667 |
| GO:0039633 | killing by virus of host cell | BP | 1 | 0.510869565 | 1 |
| GO:0044659 | viral release from host cell by cytolysis | BP | 1 | 0.512820513 | 1 |
| **GO:0003968** | RNA-directed 5'-3' RNA polymerase activity | MF | 0.480903475 | 0.019437661 | 0.346347559 |
| **GO:0047661** | amino-acid racemase activity | MF | 0.642857143 | 0.2 | 0.093073593 |
| **GO:0008238** | exopeptidase activity | MF | 0.754598846 | 0.332999142 | 0.831611136 |
| **GO:0019030** | icosahedral viral capsid | CC | 0.505780347 | 0.091666667 | 1 |
| **GO:0008237** | metallopeptidase activity | MF | 0.730667377 | 0.347115767 | 0.768600405 |
| **GO:0010181** | FMN binding | MF | 0.543060779 | 0.168609832 | 0.531734474 |
| **GO:0034061** | DNA polymerase activity | MF | 0.503498652 | 0.132337175 | 0.416682176 |
| **GO:0004180** | carboxypeptidase activity | MF | 0.657264616 | 0.308982453 | 0.730095663 |
| **GO:0097747** | RNA polymerase activity | MF | 0.523292297 | 0.178760823 | 0.612304907 |
| **GO:0034062** | 5'-3' RNA polymerase activity | MF | 0.528279096 | 0.184061032 | 0.613787522 |
| **GO:0008242** | omega peptidase activity | MF | 0.839099489 | 0.499642491 | 0.772809194 |
| **GO:0008235** | metalloexopeptidase activity | MF | 0.798551552 | 0.469072297 | 0.83554996 |
| **GO:0036459** | thiol-dependent ubiquitinyl hydrolase activity | MF | 0.900445144 | 0.573133386 | 0.801567364 |
| **GO:0005747** | mitochondrial respiratory chain complex I | CC | 0.801212998 | 0.48022218 | 0.983333333 |

**Table S2.** AUPRs of GO terms with better predictability from structure-only over sequence-only model.

| GO | GO term | task | Sequence-only | Structure-only | Sequence+Structure |
| --- | --- | --- | --- | --- | --- |
| **GO:0019031** | viral envelope | CC | 0.000350018 | 1 | 0.1 |
| **GO:0005839** | proteasome core complex | CC | 0.080430341 | 0.996732026 | 0.987272102 |
| **GO:0009538** | photosystem I reaction center | CC | 0.1 | 1 | 1 |
| **GO:0015671** | oxygen transport | BP | 0.008422677 | 0.874088713 | 0.907005539 |
| **GO:0005833** | hemoglobin complex | CC | 0.032878991 | 0.875392465 | 0.878095975 |
| **GO:0006662** | glycerol ether metabolic process | BP | 0.037785574 | 0.833956562 | 0.834029301 |
| **GO:0010499** | proteasomal ubiquitin-independent protein catabolic process | BP | 0.014935254 | 0.80367336 | 0.858815872 |
| **GO:0034987** | immunoglobulin receptor binding | MF | 0.018421424 | 0.805555556 | 0.75 |
| **GO:0018904** | ether metabolic process | BP | 0.03259338 | 0.764446168 | 0.784910714 |
| **GO:0030288** | outer membrane-bounded periplasmic space | CC | 0.037568365 | 0.746115405 | 0.834994734 |
| **GO:0046940** | nucleoside monophosphate phosphorylation | BP | 0.140335496 | 0.836811004 | 0.843889519 |
| **GO:0019877** | diaminopimelate biosynthetic process | BP | 0.185185185 | 0.833333333 | 0.392857143 |
| **GO:0042611** | MHC protein complex | CC | 0.236180853 | 0.868707483 | 0.961734694 |
| **GO:0042597** | periplasmic space | CC | 0.044804497 | 0.665662494 | 0.829091316 |
| **GO:0043190** | ATP-binding cassette (ABC) transporter complex | CC | 0.029957143 | 0.633781889 | 0.274275519 |
| **GO:0016833** | oxo-acid-lyase activity | MF | 0.134657277 | 0.731448413 | 0.717261905 |
| **GO:0034219** | carbohydrate transmembrane transport | BP | 0.0416284 | 0.608886678 | 0.64168028 |
| **GO:0098533** | ATPase dependent transmembrane transport complex | CC | 0.037442034 | 0.603484875 | 0.22486509 |
| **GO:0015144** | carbohydrate transmembrane transporter activity | MF | 0.257598442 | 0.821706994 | 0.891284132 |
| **GO:0010257** | NADH dehydrogenase complex assembly | BP | 0.02067914 | 0.584309815 | 0.850965406 |
| **GO:0032981** | mitochondrial respiratory chain complex I assembly | BP | 0.021210449 | 0.578296893 | 0.850598099 |
| **GO:0030313** | cell envelope | CC | 0.099814634 | 0.651919957 | 0.874100818 |
| **GO:0015669** | gas transport | BP | 0.020696337 | 0.559137371 | 0.587183775 |
| **GO:0008643** | carbohydrate transport | BP | 0.047395237 | 0.573559466 | 0.689976611 |
| **GO:0008218** | bioluminescence | BP | 0.000827713 | 0.501315789 | 0.125773994 |

**Table S3.** AUPRs of GO terms with better predictability from sequence-only over LM-GVP model.

| GO | GO term | task | Sequence-only | Structure-only | Sequence+Structure |
| --- | --- | --- | --- | --- | --- |
| **GO:0016855** | racemase and epimerase activity, acting on amino acids and derivatives | MF | 0.833333333 | 0.152631579 | 0.071794872 |
| **GO:0036361** | racemase activity, acting on amino acids and derivatives | MF | 0.75 | 0.233333333 | 0.069411765 |
| **GO:0047661** | amino-acid racemase activity | MF | 0.642857143 | 0.2 | 0.093073593 |
| **GO:0099094** | ligand-gated cation channel activity | MF | 0.611661857 | 0.344005535 | 0.406452375 |
| **GO:0003796** | lysozyme activity | MF | 1 | 0.505256648 | 0.854166667 |
| **GO:0003968** | RNA-directed 5'-3' RNA polymerase activity | MF | 0.480903475 | 0.019437661 | 0.346347559 |
| **GO:0030682** | mitigation of host defenses by symbiont | BP | 0.254083622 | 0.017744474 | 0.12784732 |
| **GO:0018024** | histone-lysine N-methyltransferase activity | MF | 0.551334781 | 0.412385246 | 0.443394616 |
| **GO:0042178** | xenobiotic catabolic process | BP | 0.13221182 | 0.008117095 | 0.029824954 |
| **GO:0046173** | polyol biosynthetic process | BP | 0.173973669 | 0.014808723 | 0.073239294 |
| **GO:0036459** | thiol-dependent ubiquitinyl hydrolase activity | MF | 0.900445144 | 0.573133386 | 0.801567364 |
| **GO:0016894** | endonuclease activity, active with either ribo- or deoxyribonucleic acids and producing 3'-phosphomonoesters | MF | 0.437391304 | 0.208403674 | 0.343305286 |
| **GO:0034061** | DNA polymerase activity | MF | 0.503498652 | 0.132337175 | 0.416682176 |
| **GO:0003684** | damaged DNA binding | MF | 0.261815821 | 0.059978214 | 0.180120532 |
| **GO:0016846** | carbon-sulfur lyase activity | MF | 0.464867425 | 0.357905779 | 0.385106564 |
| **GO:0016829** | lyase activity | MF | 0.637267563 | 0.408398336 | 0.560307538 |
| **GO:0101005** | ubiquitinyl hydrolase activity | MF | 0.9061601 | 0.669037003 | 0.834046481 |
| **GO:0003774** | motor activity | MF | 0.262622343 | 0.006011441 | 0.191448181 |
| **GO:0003887** | DNA-directed DNA polymerase activity | MF | 0.461435822 | 0.214752411 | 0.394313091 |
| **GO:0004550** | nucleoside diphosphate kinase activity | MF | 0.924569189 | 0.91808021 | 0.857487674 |
| **GO:0008242** | omega peptidase activity | MF | 0.839099489 | 0.499642491 | 0.772809194 |
| **GO:0016668** | oxidoreductase activity, acting on a sulfur group of donors, NAD(P) as acceptor | MF | 0.671214877 | 0.683386425 | 0.61191512 |
| **GO:0006586** | indolalkylamine metabolic process | BP | 0.082020265 | 0.013229232 | 0.024697572 |
| **GO:0046209** | nitric oxide metabolic process | BP | 0.090468994 | 0.022600092 | 0.034714297 |
| **GO:0016830** | carbon-carbon lyase activity | MF | 0.495853967 | 0.412075369 | 0.440608859 |

**Table S4.**
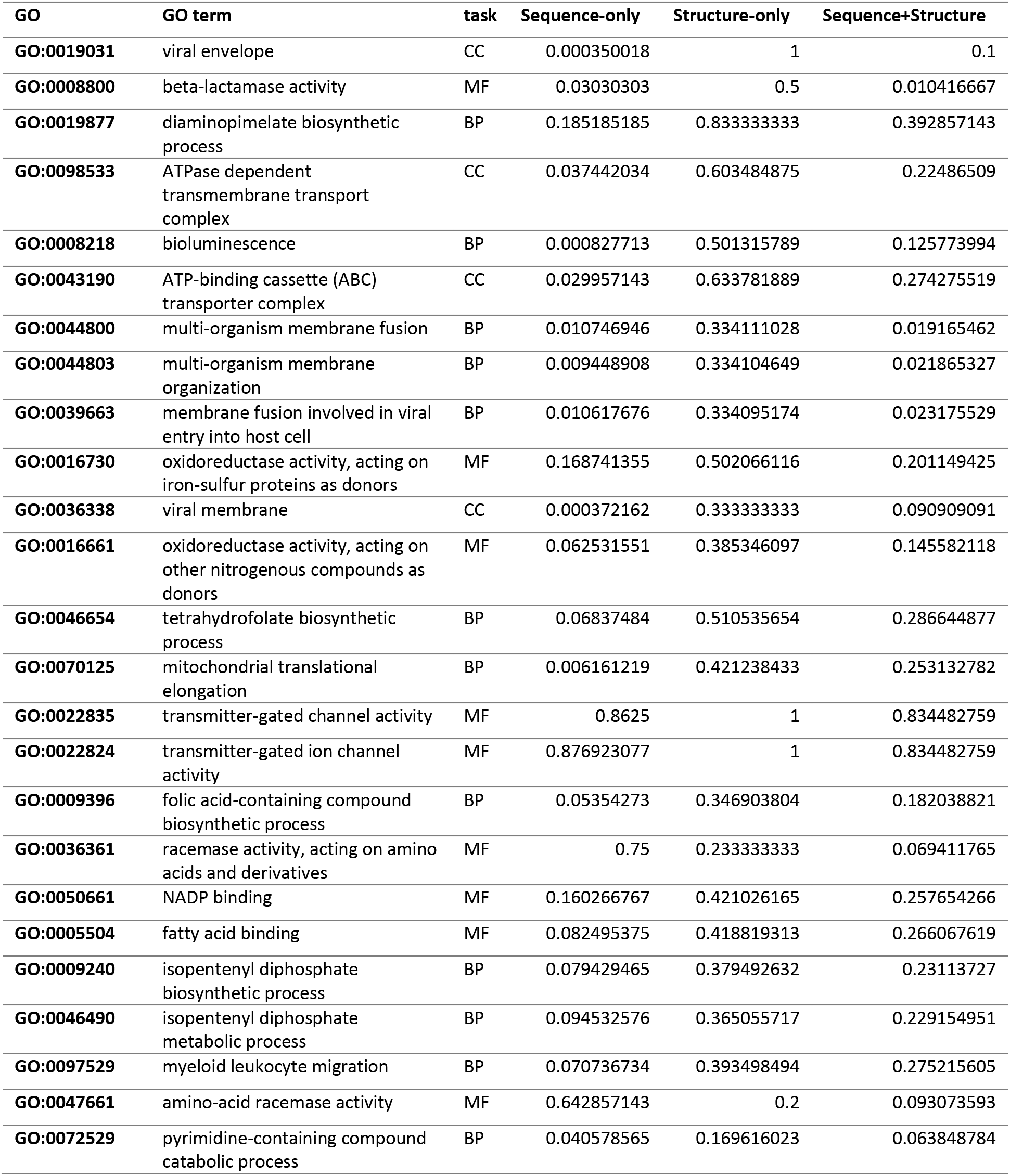
AUPRs of GO terms with better predictability from structure-only over LM-GVP model.

**Table S5.** AUROC quantifying the agreement between saliency scores from LM-GVP and <u>known active sites responsible for</u> respective MF.

| **Protein** | **GO-MF** | **AUROC** |
| --- | --- | --- |
| **1ZBD-A** | GTP binding | 0.63258 |
| **4ARZ-A** | GTP binding | 0.83401 |
| **2WKQ-A** | GTP binding | 0.87747 |
| **1J2J-A** | GTP binding | 0.66708 |
| **3RAP-R** | GTP binding | 0.68174 |
| **2A5F-A** | GTP binding | 0.79464 |
| **1E2Q-A** | ATP binding | 0.78003 |
| **1UA2-A** | ATP binding | 0.81737 |
| **3MN5-A** | ATP binding | 0.68845 |
| **2QXL-A** | ATP binding | 0.73843 |
| **1C0F-A** | ATP binding | 0.74984 |
| **2PAA-A** | ATP binding | 0.53931 |
| **4AAR-A** | ATP binding | 0.76359 |
| **2RCM-A** | Heme binding | 0.73864 |
| **1KQG-C** | Heme binding | 0.67094 |
| **1SY7-A** | Heme binding | 0.79512 |
| **2W0A-A** | Heme binding | 0.70675 |
| **1A4E-A** | Heme binding | 0.78301 |
| **2IAG-A** | Heme binding | 0.74512 |
| **1A9W-E** | Heme binding | 0.62762 |
| **2QRW-A** | Heme binding | 0.58025 |

